# Transcriptional census of epithelial-mesenchymal plasticity in cancer

**DOI:** 10.1101/2021.03.05.434142

**Authors:** David P. Cook, Barbara C. Vanderhyden

## Abstract

Epithelial-mesenchymal plasticity (EMP) contributes to tumour progression, promoting therapy resistance and immune cell evasion. Definitive molecular features of this plasticity have largely remained elusive due to the limited scale of most studies. Leveraging scRNA-seq data from 160 tumours spanning 8 different cancer types, we identify expression patterns associated with intratumoural EMP. Integrative analysis of these programs confirmed a high degree of diversity among tumours. These diverse programs are associated with combinations of various common regulatory mechanisms initiated from cues within the tumour microenvironment. We highlight that inferring regulatory features can inform effective therapeutics to restrict EMP.

## INTRODUCTION

Epithelial-mesenchymal plasticity (EMP) refers to the ability of cells to interconvert between epithelial and mesenchymal phenotypes, dynamically adopting mixed features of these states in response to signals in the cells’ microenvironment (Yang et al., 2020). Throughout a tissue, cells with phenotypes spanning an epithelial/ mesenchymal (E/M) continuum can be observed, emerging in response to specific features of their local environment. At the leading edge of tumours, for example, epithelial architecture becomes progressively disorganized and the cancer cells express higher levels of mesenchymal-associated genes (Gabbert et al., 1985; Puram et al., 2017). In cancer, this plasticity has been broadly associated with promoting metastasis, chemoresistance, and immunosuppression (Dongre and Weinberg, 2019).

Given its supposed impact on tumour progression and treatment, understanding the molecular mechanisms that drive EMP and developing therapeutic strategies to modulate it have been a priority for years (Bhatia et al., 2020; Horn et al., 2020; Ramesh et al., 2020; Yang et al., 2020). Identifying molecular determinants of EMP has largely focused on studying dynamics associated with the epithelial-mesenchymal transition (EMT) induced in experimental settings, through the addition of exogenous cytokines (eg. TGFB1) or genetic manipulation. Over the last two decades, however, it has become increasingly clear that molecular features of the EMT are highly context specific (Cook and Vanderhyden, 2020; Peixoto et al., 2019; Stemmler et al., 2019; Taube et al., 2010; Williams et al., 2019; Yang et al., 2020). The reliability of even the most canonical EMP genes (eg. SNAI1, SNAI2, CDH1, CDH2, VIM) has become unclear and the reliance on specific genes as molecular markers of EMP has led to controversy about its implication in metastasis (Aiello et al., 2017; Fischer et al., 2015, 2017; Ye et al., 2017; Zheng et al., 2015). As a result, recent guidelines from “the EMT International Association” suggest that the primary criteria for defining EMP should focus on changes to cellular properties (eg. loss of cell-cell junctions, enhanced migratory capacity) (Yang et al., 2020). Given the increasing number of genes becoming associated with EMP in the literature, this recommendation will be helpful in avoiding erroneous conclusions based on the expression of a small number of genes. In certain settings, however, assessing cellular properties is not particularly reasonable. Many studies depend on retrospective analysis of samples (eg. data generated from tumour samples) and it may not be feasible to faithfully recapitulate putative EMP phenomena ex vivo. Comprehensive interrogation into the molecular properties of EMP and, importantly, the diversity of features across contexts is critical to enable reliable interpretation of data from samples that are not amenable to phenotypic assessment. Also, embracing the diversity of EMP gene expression programs will also allow for an updated conceptual model that may help explain its involvement in tumour progression and how it may be addressed therapeutically.

The advent of single-cell transcriptomics (scRNA-seq) has enabled the identification of coordinated gene expression patterns within individual cells. Developments to increase the throughput of these assays have allowed for the parallel measurement of gene expression in thousands of single cells, sampling the phenotypic diversity among a population within a tissue (Klein et al., 2015; Macosko et al., 2015). Under the assumption that intratumoural EMP is reasonably prevalent, single-cell genomics holds the promise of revealing its intrinsic molecular characteristics. Supporting this assumption, studies applying these methods have independently identified heterogeneous expression of EMT associated genes in a variety of cancers, including squamous cell carcinoma (SCC) (Ji et al., 2020; Puram et al., 2017; Sharma et al., 2018), pancreatic (PDAC) (Raghavan et al. 2020), colorectal (Ganesh et al., 2020), and lung (Laughney et al., 2020). Detailed exploration of these programs has been limited, however, and little has been done to compare features of these programs between individual tumours and cancers. Kinker *et al*. (Kinker et al., 2020) recently assessed sources of heterogeneity in the expression profiles of monolayer-cultured cancer cell lines and identified three recurrent programs consistent with EMP, but, the similarity of these programs to those that emerge in solid tumours is unclear.

Here, we leveraged scRNA-seq data from 160 tumour samples spanning 8 cancer types to identify coordinated expression programs consistent with intratumoural EMP. While the overall composition of these programs was highly variable, we derived a set of well-conserved genes from these programs that can serve as a general EMP signature. We used this signature to query the pan-cancer data from The Cancer Genome Atlas (TCGA) and found that EMP was associated with reduced progression-free intervals and changes in immune cell proportions within the tumour microenvironment (TME). Inferences of regulatory mechanisms contributing to these EMP programs suggest that the diversity of these programs can arise from common underlying regulatory mechanisms, including ubiquitous activation of TGFB1/NFkB/TNF signalling, but that some programs have notably high MAPK or STAT/hypoxia signalling activity. We integrate transcriptomic data from kinase inhibitor screens to demonstrate that these common regulatory mechanisms present promising targets for therapeutic restriction of EMP.

## RESULTS

### A pan-cancer census of EMT-associated gene expression

We first collected droplet-based scRNA-seq data from 12 studies of 8 different cancer types, including colorectal (Lee et al., 2020; Qian et al., 2020; Uhlitz et al.), gastric (Sathe et al., 2020), lung (Kim et al., 2020; Lambrechts et al., 2018; Laughney et al., 2020; Qian et al., 2020), uveal melanoma (Durante et al., 2020), squamous cell(Ji et al., 2020), ovarian (Geistlinger et al., 2020; Qian et al., 2020), pancreatic (Steele et al., 2020), and breast (Qian et al., 2020). After removing samples with fewer than 100 malignant cells, the data comprises expression profiles of 182,198 cancer cells from 160 tumour samples (**Figure 1A; Figure S1**).

**Figure 1.**
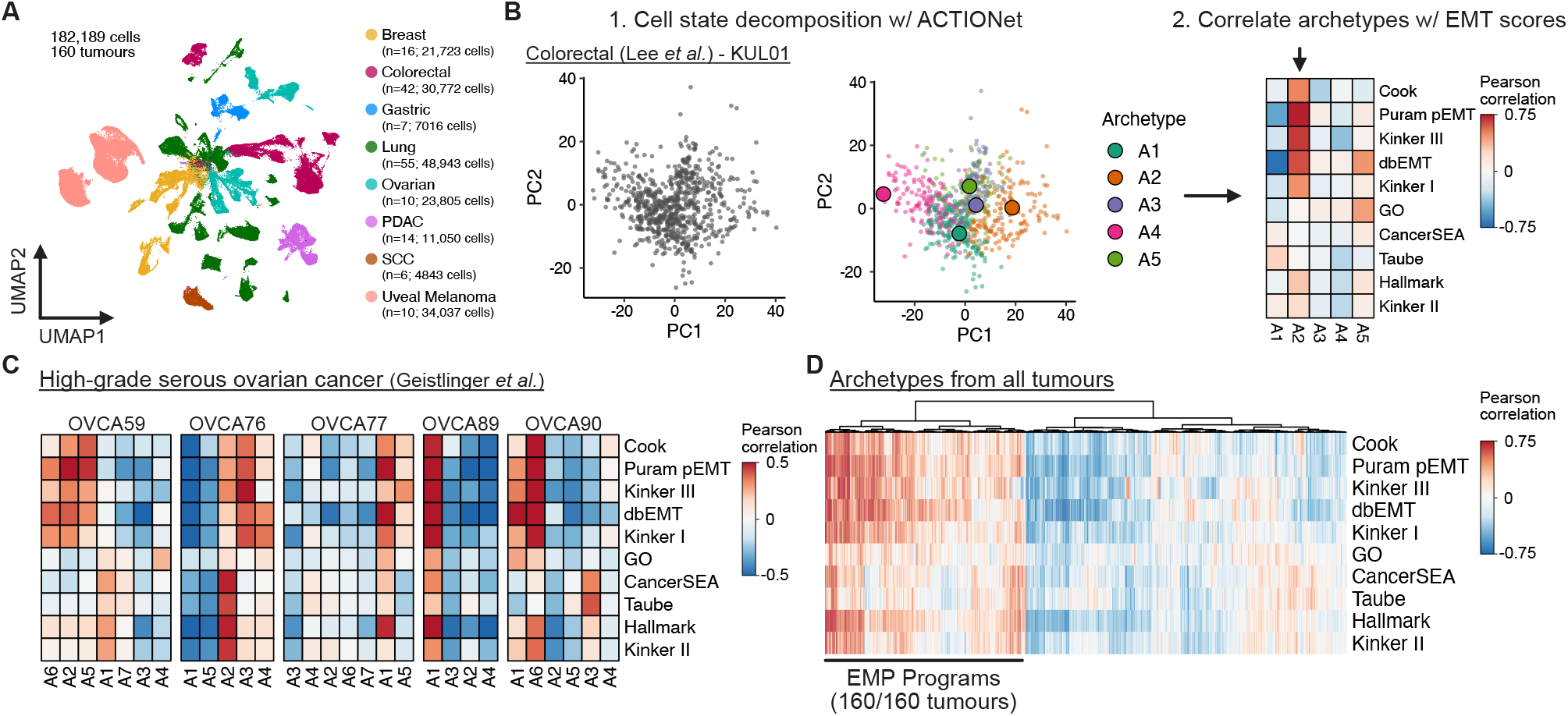
Using archetypal analysis to learn EMP-associated gene expression programs. **(A)** UMAP embedding of malignant cell scRNA-seq data from 160 tumours and 12 different studies. **(B)** Schematic representation of the analysis strategy to identify sample-specific EMP programs using archetypal analysis and correlating archetype scores with EMT gene set scores. **(C)** Pearson correlation coefficients of archetype scores with EMT gene set scores of individual cells from 5 high-grade serous ovarian tumours. **(D)** Hierarchically clustered heatmap of Pearson correlation coefficients of all archetype scores with EMT gene set scores from the 160 tumours analyzed.

To assess intratumoural heterogeneity of EMT-associated gene expression, we scored each cell for its relative expression of genes contained in 10 different EMT gene sets, including several curated sets [the “Epithelial-mesenchymal transition” GO Term, MSigDB Hallmark gene set (Liberzon et al., 2015), CancerSEA (Yuan et al., 2019), dbEMT1.0 (Zhao et al., 2015)] and others derived from individual studies [Taube (Taube et al., 2010), Puram (Puram et al., 2017), Cook (Cook and Vanderhyden, 2020), and Kinker I, II, III (Kinker et al., 2020)]. The composition of these gene sets is high variable, with over two-thirds of the genes being present in only a single gene set (**Figure S2A**). While it is possible these diverse gene sets could represent different subsets of a common underlying biological process, we correlated the gene set scores across all cells and found that this is not uniformly true. The 10 gene sets formed two groups of well-correlated scores, suggesting that they may represent distinct EMP programs (**Figure S2B**).

Gene set scores can provide biological insight into gene expression patterns, but they can be influenced by uninteresting features of the data. Specifically, they can be inflated by high expression of a small proportion of the set’s genes that are not necessarily determinants of the queried biological process. Variation in scores across a population of cells may also reflect random fluctuations of the set’s genes and not necessarily a coordinated activation of the process. Matrix factorization approaches have been applied to scRNA-seq data to learn coordinated expression programs heterogeneously expressed across a population. By learning these programs from the data itself, reliance on previously defined gene sets is restricted to only the interpretation of the programs.

We next sought to explore heterogeneously expressed programs from the 160 tumours. To identify these latent gene expression programs, we performed multi-resolution archetypal analysis on each tumour sample using the ACTIONet algorithm (Mohammadi et al., 2020), learning cell activity scores for distinct programs in each tumour (**Figure 1B**). This identified multiple programs in each sample, including expected sources of variation, such as cell cycle activity. To identify those associated with EMP, we correlated the cellular activity of each latent program learned by ACTIONet with the previous EMT gene set scores. All 160 tumours had latent programs well-correlated with these scores, suggesting intratumoural EMP is a ubiquitous feature of solid tumours and readily captured in scRNA-seq experiments (**Figure 1C**,**D**). Kinker et al. (**Kinker et al**., **2020**) recently demonstrated recurrent heterogeneity of programs consistent with EMP in scRNA-seq data derived from monolayer cultures of cancer cell lines, but this is the first quantitative demonstration of this plasticity in solid tumours across multiple cancer types.

### Defining a conserved signature associated with EMP

To define the specific genes contributing to EMP in each tumour, we identified genes whose expression significantly changes as a function of the cells’ activity of each latent program learned from ACTIONet. We have previously shown that transcriptional responses of experimentally induced EMTs are highly context-specific, but it was unclear if the same diversity existed in vivo (Cook and Vanderhyden, 2020). Of the 3822 genes differentially expressed in at least one EMP program, the vast majority were associated with a small number of samples. We clustered differentially expressed genes based on their model coefficients for each EMP program and identified a group of 640 genes with a frequent positive association (**Figure 2A**). We further refined this signature by removing genes that were associated with fewer than 10 EMP programs or also downregulated in more than 10 EMP programs, resulting in an EMP signature of 289 genes (**Figure 2B**).

**Figure 2.**
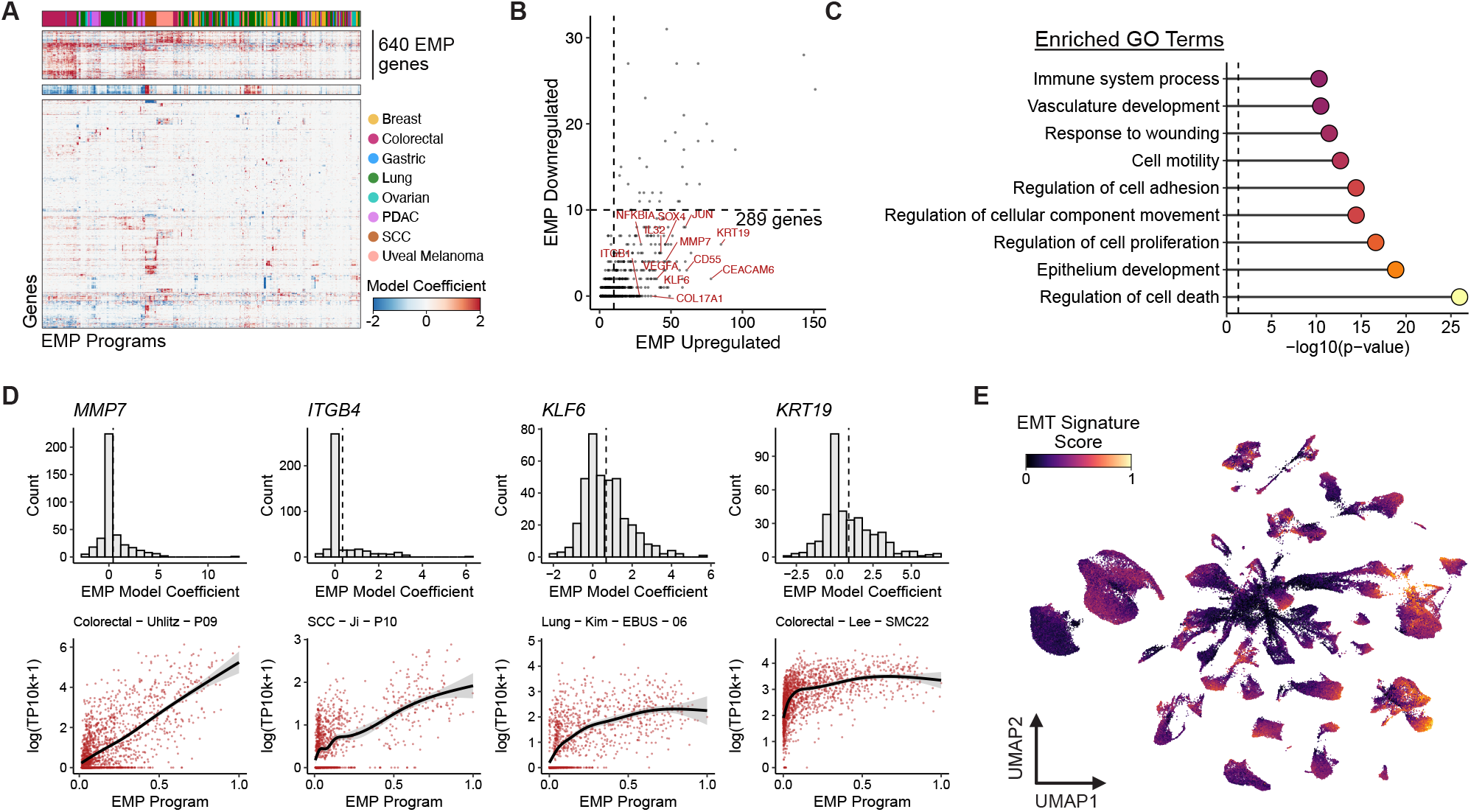
Defining a conserved EMP gene signature. **(A)** Hierarchically clustered heatmap of EMP model coefficients for 3822 genes differentially expressed in at least one EMP program. **(B)** Plot showing how frequently each of the 640 genes were associated with down- and upregulation in EMP programs. **(C)** EMP-associated GO terms significantly enriched in the 289 conserved EMP genes. P-values were calculated using Fisher’s exact tests and adjusted using the Benjamini-Hochberg method. **(D)** Examples of EMP associated genes. Top plots show the distribution of effect sizes for each gene with all EMP programs. Dashed line represents the mean value. Bottom plots show expression values (log-transformed UMI counts per 10,000 UMIs) for each gene in the EMP program with the highest effect size for that gene. **(E)** UMAP embedding of malignant cells from the 160 tumours analyzed, coloured by a gene set score for the signature of 289 EMP-associated genes.

EMP programs were associated with reduced expression of a group of 125 genes. Interestingly, epithelial genes were not enriched among this group. Rather, they strongly enrich for GO terms associated with cell cycle (p = 2.9e-73, Fisher’s exact test). This is consistent with the proliferation-migration trade-off associated with a mesenchymal phenotype. We do note that several programs classified as EMP programs in multiple tumour types showed activation of these genes. The lack of expression patterns associated with loss of an epithelial phenotype further supports the growing evidence of hybrid E/M phenotypes being highly prevalent in cancer (Kröger et al., 2019; Pastushenko and Blanpain, 2019; Pastushenko et al., 2018; Puram et al., 2017).

Of the 289 genes positively associated with EMP, no individual gene was a perfect indicator of its activity, but as a collective, they represent a fairly consistent signature. They also enrich for GO Terms consistent with a mesenchymal phenotype, including cell motility, regulation of cell adhesion, and response to wounding (**Figure 2C**). Many of the canonical EMT genes are not included in this signature, including *CDH2, VIM, SNAI1, SNAI2*, and *ZEB1*, although these genes did have variable associations with EMP programs (**Figure S3**). The signature did, however, include many genes that have previously been implicated in the EMT, including various transcription factors (*SOX4, KLF2/4/6/10, JUND*), integrins (*ITGB1/4/8, ITGA2/3*), secreted factors (*VEGFA, IL32, CXCL1, CXCL8*), and membrane proteins (*CD24, CD59, S100A6, CEACAM1*), and more (**Figure 2D, Supplemental Data**).

The stability and distribution of phenotypes along an epithelial-mesenchymal continuum has gained attention recently, with the relevance of hybrid phenotypes being contrasted to fully epithelial or mesenchymal cells (Aiello et al., 2018). Using gene set scores of the conserved signature as a relative measure of the cells’ mesenchymal activity, we found that different tumours can have different average levels of mesenchymal expression, but also that the variance of scores can vary dramatically, suggesting that range of phenotypes represented within the tumour can vary (**Figure 2E; Figure S4**). We also note that the majority of tumours don’t have clear multimodal distributions that would be consistent with a model where various states along the phenotypic continuum have elevated stability. Rather, cells span the continuum, forming a distribution with most cells occupying intermediate states and tails spreading to more extreme phenotypes. These patterns are recapitulated when using gene set scores from the EMT-associated gene sets explored previously and the specific EMP-associated program learned from the tumours’ themselves (**Figure S4**).

### A refined, malignant-cell specific EMP signature is associated with poor patient prognosis

Due to their inherent similarities, the ability to distinguish fibroblast and EMP-specific expression patterns has been a challenge. Many EMT gene sets contain genes highly expressed in fibroblast populations and as a result, “mesenchymal” features of tumours defined from bulk RNA-seq data have been found to often be associated with fibroblast content of the tumour rather than cancer cell plasticity (Isella et al., 2015; Izar et al., 2020; Puram et al., 2017). The choice of specific markers used to assess EMP in studies has also led to controversy (Yang et al., 2020). This confusion has made it challenging to draw conclusions about the involvement of EMP in tumour progression and clinical outcomes.

For each tumour sample, we calculated a cell type specificity score for each of the 289 conserved EMP genes and averaged these scores across tumours to get an overview of how specific the markers were to cancer cells (**Figure 3A**). Of the 289 genes, 165 were highly specific to cancer cells, whereas the remaining 124 were also expressed in fibroblasts, macrophages, and/or T cells. A signature of cancer cell-specific EMP genes could be valuable for generating EMP activity scores in scRNA-seq data, so we established a refined signature comprising the 165 genes highly specific to cancer cells (**Supplemental Data**). We scored individual cells comprising 22 colorectal tumours(Lee et al., 2020) for their activity of this signature and found heterogeneous activity among cancer cell populations, with effectively no activity in non-malignant cell types (**Figure 3B**).

**Figure 3.**
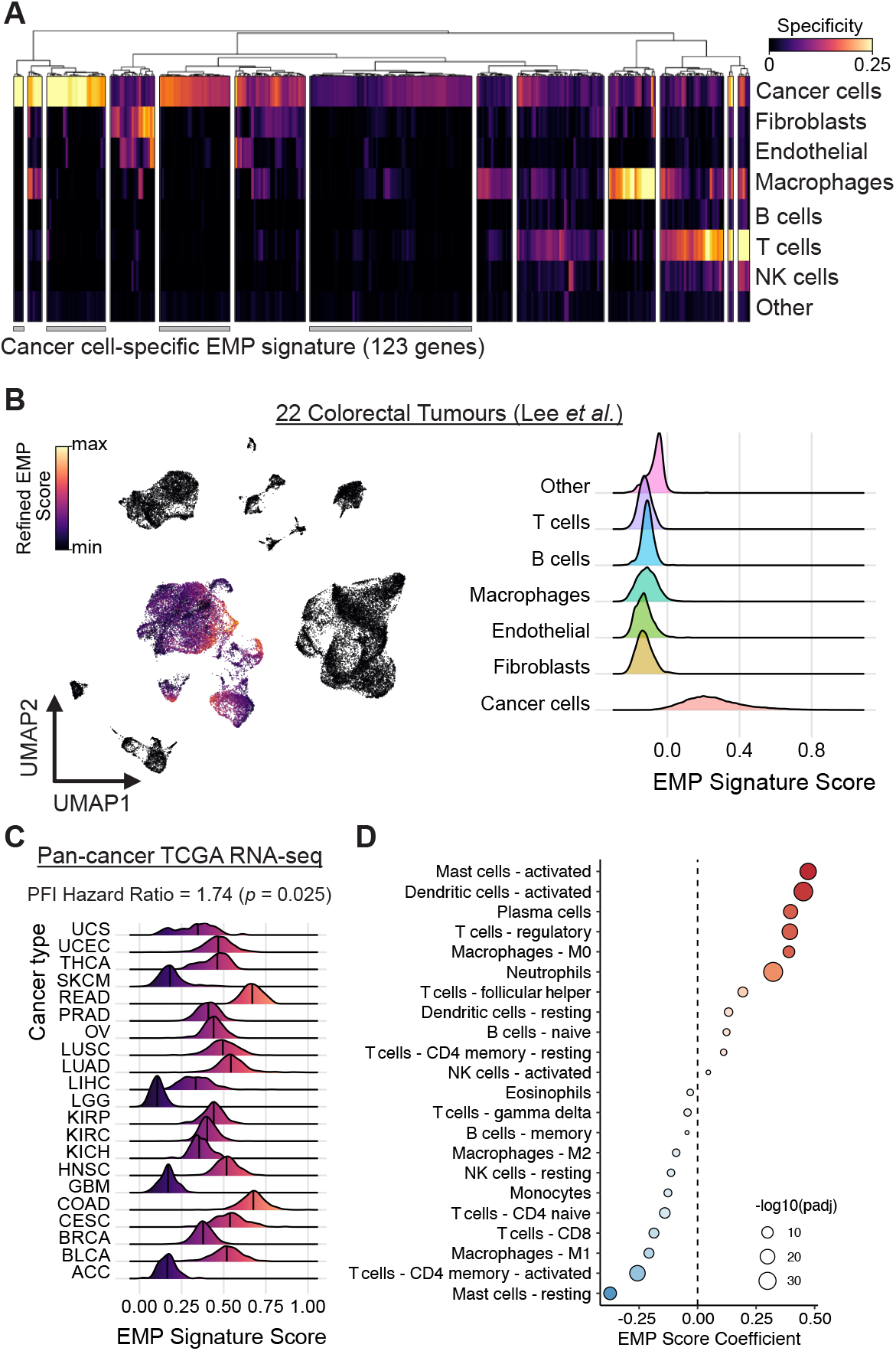
EMP is associated with worse progression-free survival and an immunosuppressive TME. **(A)** Clustered heatmap of cell type specificity scores for each of the 289 EMP-associated genes. Clusters of genes with high cancer cell specificity were defined as a cancer cell-specific EMP signature. **(B)** UMAP embedding (left) and density plot (right) showing the distribution of gene set scores for the cancer cell-specific EMP signature in all cell types from 22 colorectal tumours. **(C)** Progression-free interval (PFI) hazard ratio and distribution of signature scores from the TCGA’s pan-cancer bulk RNA-seq cohort. **(D)** Changes in immune cell proportion estimates as a function of EMP signature scores from the TCGA’s pan-cancer cohort.

Given this high specificity, the signature could be used as a measure of mesenchymal properties in bulk RNA-seq data without being confounded by expression from fibroblasts. We used the pan-cancer RNA-seq data from The Cancer Genome Atlas (TCGA) (Hoadley et al., 2018) to calculate a relative EMP score for all tumours. Modelling patients’ progression-free interval (PFI) as a function of this signature activity, tumour type, age at diagnosis, and tumour purity, we found that EMP activity was associated with a reduced PFI (Cox hazard ratio: 1.74; p = 0.025) (**Figure 3C**). Using estimates of immune cell proportions across all TCGA samples (Thorsson et al., 2019), we also found high expression of this mesenchymal signature was associated with larger proportions of inflammatory cell types (activated mast and dendritic cells), but also more immunosuppressive regulatory T cells and fewer effector cell types, including CD8 T cells and naive CD4 T cells (**Figure 3D**).

### Diverse paracrine signalling modulates EMP

While unifying molecular signatures are appealing, appreciating the diversity of EMP programs is critical as it may contribute to functional nuances of the phenotype. These programs may also have varying regulatory dependencies that would warrant different therapeutic approaches. Variation in cell state emerges from complex differences in the cells’ microenvironment, including changes in oxygen concentration, nutrient availability, cytokine and growth factor levels, and juxtacrine interactions with adjacent cell types. These factors converge on signal transduction pathways and transcription factor networks, whose output is further tuned by epigenetic and mutational features of the cell. We next sought to use computational approaches to infer how regulation at these levels contribute to the patient-specific EMP programs we learned from scRNA-seq data.

We used PROGENy’s signalling activity model (Holland et al., 2020a, 2020b; Schubert et al., 2018) to calculate relative activity scores for 14 signalling pathways and assessed their activity as a function of EMP program activity in each tumour sample. EMP programs were consistently associated with elevated TGFB, NFkB, and TNFa signalling, but some had distinctly high levels of either EGFR and MAPK signalling or JAK-STAT, Hypoxia, and p53 signalling (**Figure 4A**). This suggests that hypoxia-associated EMP may have distinct features from plasticity coordinated by MAPK/ERK signalling.

**Figure 4.**
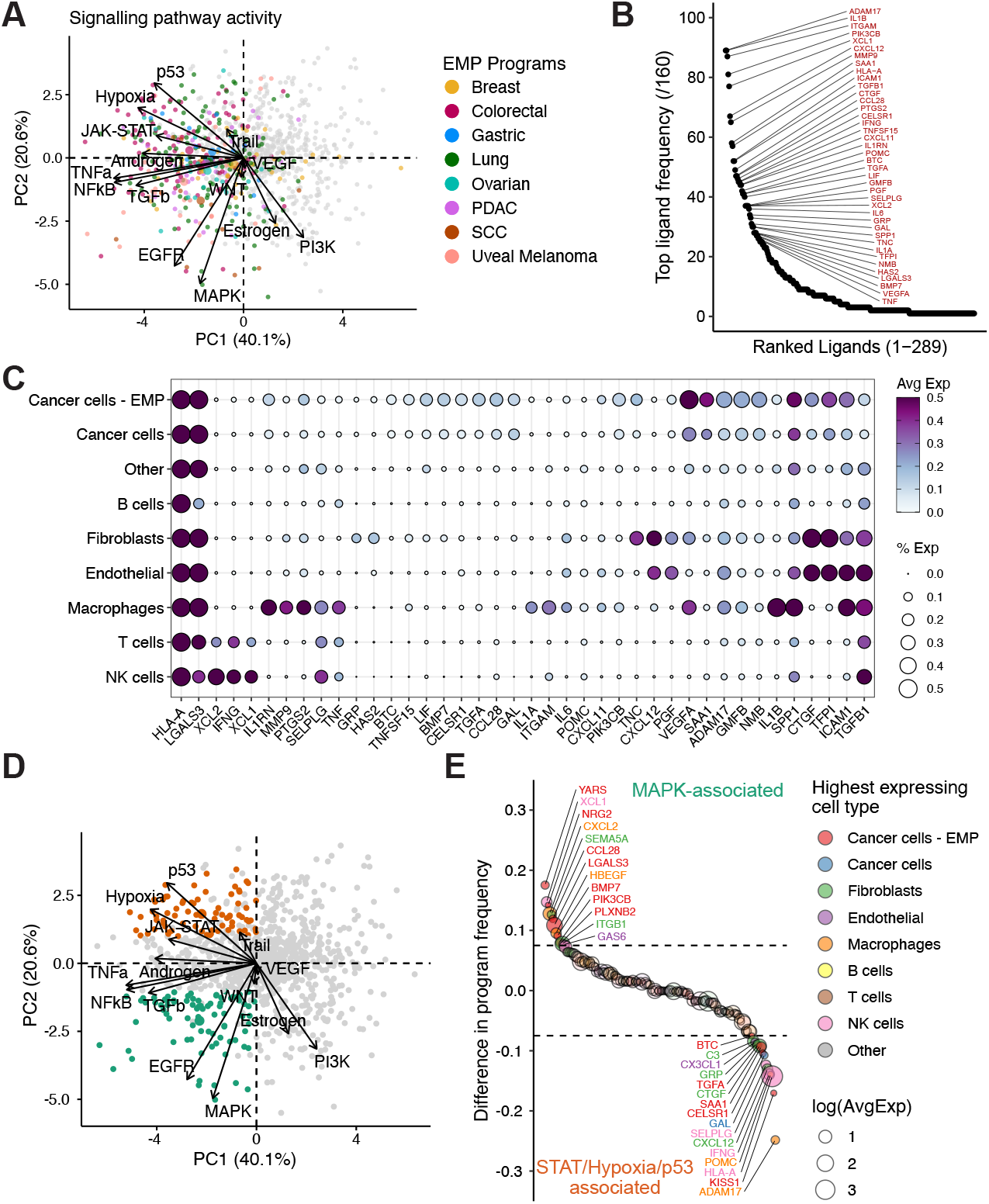
Regulatory mechanisms contributing to EMP expression programs. **(A)** PCA of model coefficients for PROGENy’s signalling pathway activity inferences for all latent programs. Grey dots represent programs not associated with EMP. **(B)** Counts for the number of EMP programs (/160) in which each ligand was implicated as a top regulator. Ligands implicated in 30 or more programs are highlighted. **(C)** Expression of top ligands in cell types throughout the tumour microenvironment. Values are calculated as averages of all 160 tumours. **(D)** Same as (A), but coloured to define STAT/Hypoxia/p53 (orange) and MAPK-associated (green) EMP programs. **(E)** Ligands inferred to preferentially contribute to MAPK- or STAT/Hypoxia/ p53-associated EMP. Values represent the difference in the proportion of programs in which each ligand was implicated.

Differences in the activity of these pathways could be related to the cells’ association with non-malignant cells of the TME, which secrete many of the factors that can promote EMP. Since the data sets we have used included matched gene expression profiles of stromal cells for all tumours, we inferred cell-cell communication for each tumour to identify ligands from the TME that could account for that sample’s specific EMP expression program. Several cytokines were associated with EMP in many tumours, including *TGFB1, IL6, IFNG, TNFSF15*, and others (**Figure 4B**). Consistent with previous literature, macrophages and fibroblasts were dominant sources of many factors contributing to EMP (Dongre and Weinberg, 2019; Puram et al., 2017)(**Figure 4C**). Mesenchymal malignant cells also express various ligands predicted to promote EMP, suggesting that they may establish self-regulatory signalling loops within the TME.

To identify factors that may lead to the differences between the EMP programs with high EGFR/MAPK activity and those with dominant JAK-STAT/Hypoxia/p53, we compared how frequently ligands were associated with each set of programs (**Figure 4D**,**E**). Interestingly, several ligands expressed most highly in mesenchymal cells themselves were preferentially implicated in MAPK-associated EMP (eg. *YARS, NRG2, CCL28, LGALS3, BMP7*), whereas EMP programs with high STAT/Hypoxia/p53 signalling were associated with ligands from stromal populations (eg. *IFNG, HLA-A, ADAM17, CXCL12*). Together, this highlights that although malignant populations may have cell-autonomous mechanisms to promote EMP, the molecular features of this plasticity can be modulated by complex interactions with stromal populations in the TME.

### Pharmacologic restriction of EMP

EMP has been implicated in both chemoresistance and immune cell evasion (Dongre and Weinberg, 2019). Given this, we predict that non-lethal restriction of EMP could be a promising therapeutic approach to sensitize tumours to orthogonal treatments and elicit immune cell killing. Diversity of EMP programs could introduce challenges for effectively preventing these cell dynamics, but the dependence of EMP on factors from the cells’ microenvironment suggests that the diversity likely arises from combinatorial effects from the relatively limited number of signal transduction pathways. Therefore, we suspect that diverse EMP programs may be susceptible to common pathway perturbations and rational treatments could be devised by inferring signalling activity associated with EMP in a given tumour.

To begin to test this prediction, we first explored the MIX-Seq data set comprising scRNA-seq profiles of over 100 cancer cell lines treated with various drugs, including the MEK inhibitor Trametinib (McFarland et al., 2020). We used ACTIONet to define cell line-specific EMP programs from untreated expression profiles, identifying high-confidence EMP programs in 46 of the 99 lines we assessed (lines with >100 cells; **Figure S5A**). Many others had programs that correlated well with individual EMT gene sets and may represent EMP, but out of caution, we did not annotate them as such. We then inferred changes in signalling pathway activity associated with all latent expression programs and found that, like in the tumour samples, TGFB/NFkB/TNFa activity was consistently higher in EMP programs, but programs could be distinguished by high EGFR/MAPK or STAT/ Hypoxia/p53 signalling (**Figure 5A**).

**Figure 5.**
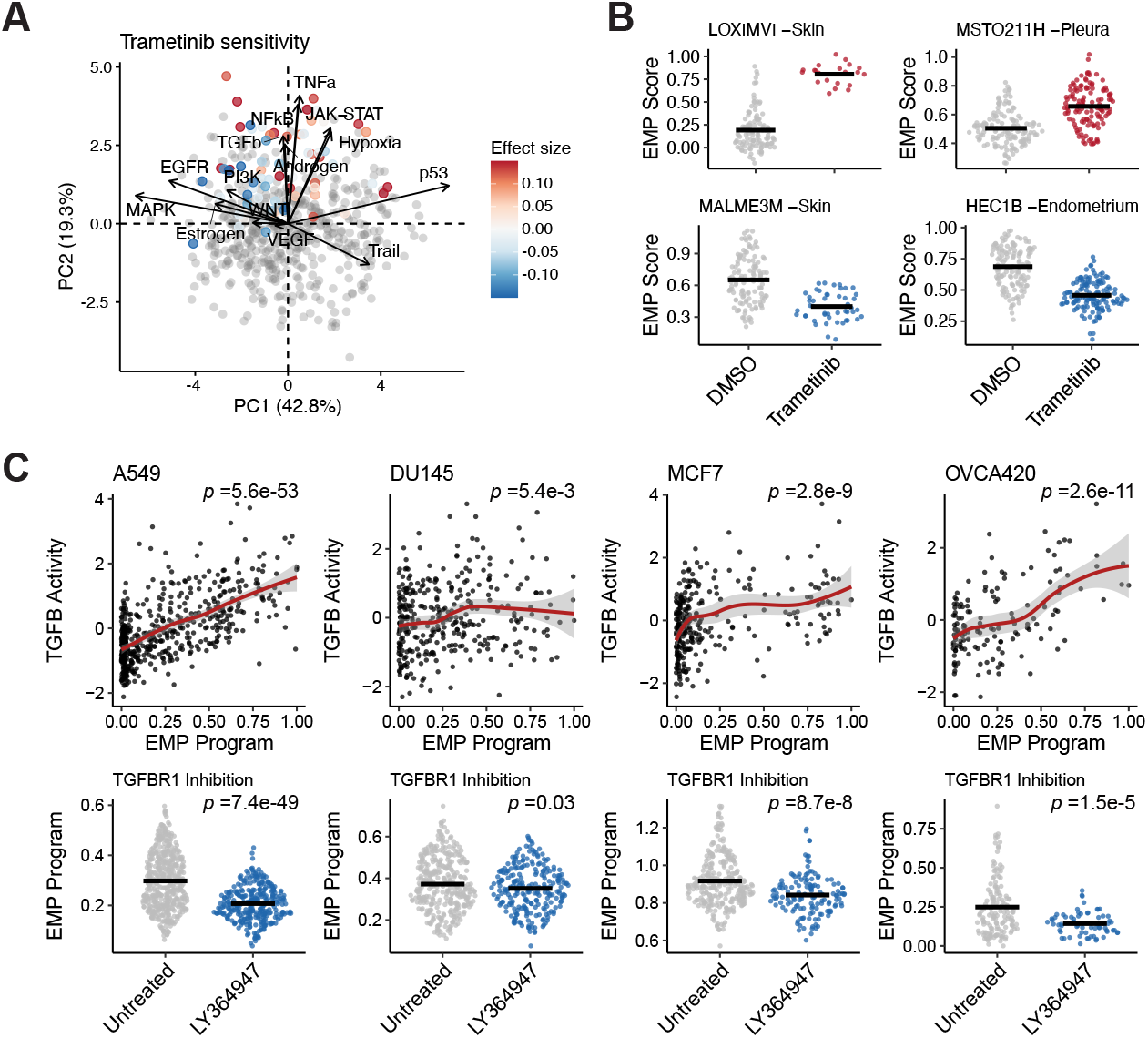
Regulatory predictions can infer strategies to therapeutically restrict EMP. **(A)** PCA of model coefficients for PROGENy’s signalling pathway activity inferences for latent programs from the 99 cell lines in the MIX-seq data set. EMP programs are coloured by the effect trametinib has on its activity. **(B)** Examples of cell lines whose specific EMP program is enhanced (top) or limited (bottom) by trametinib. **(C)** Effect of TGFBR1 inhibition by LY364947 on sample-specific EMP programs of A549, DU145, MCF7, and OVCA420 cell lines. P-values were calculated from linear models for each condition and were all corrected with the Benjamini-Hochberg method.

To determine if MEK inhibition preferentially limits MAPK-associated EMP, we used the MIX-Seq dataset to assess the effects of trametinib on all EMP programs. Trametinib had diverse effects across cell lines and even increased EMP program activity in several, but in general, the programs inferred to be associated with high MAPK signalling had reduced activity following MEK inhibition (**Figure 5A**,**B**). Given the consistent association of EMP with TGFB1 and NFkB signalling, we would also predict that inhibition of these pathways could restrict EMP. While TGFB1 and NFkB inhibitors were not included in the MX-Seq data set, we have previously published scRNA-seq data of four cancer cell lines (A549, DU145, MCF7, and OVCA420) treated with the TGFBR1 inhibitor LY364947. Performing the same analysis on this data, we found that inhibition of TGFBR1 in cells cultured in control conditions led to repression of most EMP programs, proportional to the inferred level of TGFB1 activity associated with each line’s intrinsic EMP (**Figure 5C; Figure S5B**). Together, this supports that EMP programs with highly diverse molecular features may still have common dependencies and effective strategies for restricting EMP can be discovered by inferring these regulatory mechanisms.

## DISCUSSION

EMP has long been appreciated as a prominent source of intratumoural heterogeneity that ultimately promotes tumour progression and hinders effective treatment. While select molecular patterns associated with EMP have been known for decades, variability in their involvement across contexts has raised confusion and as a result, high-level conceptual models to understand the molecular basis of this plasticity have been lacking. Here, we used scRNA-seq data of 160 tumour samples to identify gene expression programs consistent with EMP, compared features of these programs between samples, inferred causal regulatory networks driving them, and explored therapeutic options to limit their activity.

EMP programs identified in each sample were consistently associated with higher expression of genes associated with mesenchymal traits (eg. motility, cellular rearrangement), yet there was no consistent reduction of epithelial genes upon activation of these programs. This supports studies demonstrating the presence of hybrid phenotypes in cancer with distinct functional traits (Kröger et al., 2019; Pastushenko and Blanpain, 2019; Pastushenko et al., 2018). It is possible that reduction of epithelial traits is achieved through internalization of epithelial membrane proteins rather than transcription repression, as has been seen with hybrid E/M phenotypes in pancreatic cancer(Aiello et al., 2018). However, the generalizability of post-translational regulation to the broader scope of phenotypically relevant epithelial proteins is still unclear. This could, however, reconcile the presence of various epithelial membrane proteins in EMP programs, perhaps acting as a feedback mechanism to counter this regulation.

We acknowledge that our ability to define EMP programs may be limited by the sparsity and sensitivity constraints of scRNA-seq assays. It is very possible that some phenotypically relevant expression patterns were not reliably detected in our analysis for technical reasons. For example, we found that some of the canonical EMT transcription factors (eg. *SNAI1, SNAI2, ZEB1*) were poorly detected and it is unclear if this is a feature of these samples or a technical limitation. Also, given the relevance of post-transcriptional regulation of EMP (Aiello et al., 2018), it is feasible that some of the most conserved features of EMP reside beyond cells’ expression profiles. Advances in morphological and protein-based assays (eg. high-content imaging, imaging mass cytometry, etc) will allow for the discovery of additional features of EMP.

A general observation worth noting is that the E/M phenotypes observed within a single sample seem to span a continuous, often monomodal distribution. There has been much speculation about the stability of phenotypes along an E/M continuum and the presence of discrete stable states (Nieto et al., 2016). Modelling frameworks suggest their existence (Jia et al., 2017; Zadran et al., 2014), but it is not clear that they are compatible with the distributions of phenotypes observed in these samples: the dominant E/M state between tumours is variable, the extent of phenotypic variation within individual samples is highly variable, and very few samples show the multimodality that should coincide with varying stability along a phenotypic continuum. Though, considering these findings, it is intriguing to question if it is more relevant to a tumour’s biology that its malignant cells are more mesenchymal on average, or if they span a wider range of E/M phenotypes.

We identified a signature of 289 genes that were commonly associated with EMP programs and a refined signature of 165 genes with high specificity to malignant cells, which could be useful for quantifying EMP in gene expression data, as we did here with TCGA data. In no way do we argue that this signature represents the most biologically relevant components of EMP gene expression programs. Enrichment of GO terms related to mesenchymal functions suggests that they likely contribute to the phenotype, but we found a total of 3822 genes associated with EMP across the various samples. Even under highly controlled experimental conditions, EMT responses are vastly context-specific (Cook and Vanderhyden, 2020; McFaline-Figueroa et al., 2019; Peixoto et al., 2019). The phenotypic changes associated with EMP are likely an emergent property of these many changes, and variation in expression programs may provide phenotypic nuances that we are unaware of. As such, rather than try to reduce EMP to some consistent phenomenon, we think it is important to recognize context specificity and begin building conceptual models and experimental designs to understand its importance.

The ability to therapeutically restrict plasticity in tumours could greatly improve the efficacy of existing treatment options. One could imagine several strategies for accomplishing this: interfering with the cues that initiate plasticity, therapeutically impairing transcriptional regulatory mechanisms (eg. inhibiting histone modifying proteins), or targeting effector proteins associated with a given cell state (eg. neutralizing immunosuppressive cytokines released from mesenchymal cells). The latter may be challenging due to the diversity of EMP phenotypes, but is perhaps the most direct strategy for preventing undesirable features of mesenchymal cells. In this study, we have begun to explore strategies for blocking stimulatory pathways, restricting diverse programs with common signalling cues. We computationally inferred that EMP programs were consistently associated with TGFB and NFkB signalling, whereas some had notably high levels of either MAPK/EGFR or STAT/Hypoxia/p53 signalling. Leveraging transcriptomic data from various drug screens, we confirmed that targeting these active pathways led to reduced activity of sample-specific EMP programs despite the diversity of the programs themselves. While this diversity introduces challenges for effective therapeutics, this observation suggests that these diverse programs may have common dependencies that can be inferred from their molecular features and exploited therapeutically.

In summary, we have used scRNA-seq data from 160 tumours spanning 8 cancer types to identify molecular features associated with EMP. We find that EMP is a ubiquitous source of intratumoural heterogeneity but — consistent with previous findings — is highly context specific. We identify a cancer cell-specific signature of the most common genes positively associated with EMP and demonstrate its utility as a general-use gene set, using the TCGA pan-cancer RNA-seq data to associate EMP with worse progression-free intervals and a more immunosuppressive TME. We use computational approaches to infer regulatory features of EMP across hundreds of samples and highlight that diversity may emerge from common regulatory mechanisms that can be inferred and used to rationalize therapeutic strategies.

## METHODS

### Preparation of scRNA-seq data

A summary of scRNA-seq data sets used in this study are described in Table S1. To avoid comparing data collected from vastly different technologies, only droplet-based scRNA-seq data was used in the analysis. Raw UMI count matrices and cell metadata were collected from the various sources (**Table S1**). Several data sets included matched normal tissue samples. In these cases, we removed the normal samples and only proceeded with the tumour samples.

Initial quality control was performed independently for each data set using the R package Seurat v4.0(Hao et al., 2020). Cells with fewer than 200 detected genes were removed and only genes detected in more than 3 cells were included in the analysis. Cells with a high percentage of mitochondrial transcripts were also removed. As the distribution of mitochondrial reads can vary significantly between sample preparations, we manually defined thresholds for each data set based on a heuristic of “trimming the skew to result in an approximately normal distribution. Though thresholds were similar between datasets, ranging between 10-25%. All data sets were normalized using SCTransform(Hafemeister and Satija, 2019) and the percentage of mitochondrial reads was regressed out to remove any variation associated this feature. The data was then processed with principal component analysis (PCA), a nearest neighbor graph was generated on the first 30 PCs and the data were clustered at a fairly low resolution (FindClusters, resolution = 0.2) using the Louvain algorithm. For visualizations presented in the figures, UMAP embeddings were generated from the first 30 PCs

For the majority of data sets, we found that non-malignant cell types from different tumours within the same data set often clustered together and coarse-grained cell type annotation did not require data integration. However, for 2 of the data sets (**Table S1**), we found significant batch effects between samples. For these samples, we performed using the default integration pipeline with SCTransform normalization implemented in Seurat v4(Hao et al., 2020). Following integration, the data was re-clustered from the aligned embeddings and we found that coarse cell type annotations based on canonical markers matched the clustering well.

### Cell type annotation

Cell types were annotated largely based on the expression of canonical markers and supported by annotations provided by the original data source if provided. The identification of cancer cells was also supported by the trend that non-malignant cells typically cluster well between tumours, whereas the diversity of malignant cells causes them to frequently cluster separately from each other. Cancer cells were identified as clusters with clear expression of epithelial markers (eg. EPCAM, KRT19). For analyses including non-malignant cell types, we only aimed to achieve a coarse-grained annotation. We defined clusters associated with T cells (CD3E), B cells (MS4A1), NK cells (NKG7, CD3E negative), macrophages (CD14), Endothelial cells (CLDN5), and fibroblasts (COL1A1, ACTA2). Clusters that were not clearly positive for one of the canonical markers were annotated as “Other”.

### Identifying latent EMP expression programs with archetypal analysis

Samples with <100 annotated cancer cells were removed from the analysis. Multi-resolution archetypal analysis was performed independently on the cancer cells from all 160 tumours using ACTIONet v2.0.15(Mohammadi et al., 2020) to decompose cells’ gene expression profiles into a small set of latent expression programs that are heterogeneously expressed throughout the population. Reduced kernel matrices were first computed with the reduce.ace() function implemented in the R package ACTIONet with the parameter reduced_dim=25. Given that each population represented a single cell type, ACTIONet was then run with the k_max=6 option to reduce the maximum depth of decompositions and with min_cells_per_arch=5 to prevent archetypes driven by a small number of cells.

Resulting archetype footprints (program activities) were correlated with gene set scores for 10 EMT gene sets. Clustering of the Pearson correlation coefficients allowed us to define EMP-associated programs as clusters with high correlation values. We then used linear models to identify genes whose expression is associated with all latent programs. To only recover reliable changes, differential expression was limited to the top 2000 variable genes with a minimum detection frequency of 5% in each cancer cell population. Pearson residuals from SCTransform’s model were used for differential expression, modelling each gene’s residuals as a function of program activity and program-associated genes were defined as those with a with a Benjamini-Hochberg-corrected P-value of <0.05 and a model coefficient (effect size) of >0.5. For generating a conserved EMP signature, we first modified these thresholds slightly and only considered genes with a model coefficient of >2 in at least one EMP program.

### Gene set scoring

Gene set scoring was performed using Seurat’s AddModuleScore function using default options, which calculates the average expression of genes within a gene set relative to control genes of similar average expression levels.

### Cell type specificity scoring

Specificity scores were calculated for EMP signature genes using the R package genesorteR v0.4.3(Ibrahim and Kramann, 2019). Specificity scores represent the exclusivity of a gene to a given cluster and extent to which it is expressed (proportion of cells with detection greater than population median levels). Values range from 0-1, with 1 corresponding to genes that are exclusive to a given cluster and detected in all cells.

### Pan-cancer TCGA analysis

Bulk RNA-seq profiles and clinical outcomes were accessed from https://gdc.cancer.gov/node/905, tumor purity estimates were acquired from Aran et al.(Aran et al., 2015), and immune cell proportion estimates were accessed from Thorsson et al.(Thorsson et al., 2019).

RNA-seq count data for each sample was normalized to counts per million (CPM) and log-transformed. EMP signature scores were calculated for each sample by computing the average gene-level Z-score of the genes from the 165 cancer cell-specific genes. The association of EMP signature activity with progression free interval was assessed using a Cox proportional hazards model, including tumour type, purity, age, and sex along with continuous EMP scores as covariates in the model. Changes in immune cell proportions were assessed by independently using linear models to model each cell type’s predicted proportion as a function EMP signature score, including tumour type as a covariate. We used the Benjamini-Hochberg method to correct p-values to account for multiple comparisons.

### Inferring EMP-associated signalling activity

The R package PROGENy v1.11.2(Holland et al., 2020a, 2020b; Schubert et al., 2018) was used to infer the activity of 14 signalling pathways in each individual cell. For each tumour sample, Pearson residuals from the SCTransform model were used to calculate pathway activity with the progeny() function with the top=500 parameter set to use the top 500 genes of each pathway in PROGENy’s model for the activity calculation. We used simple linear models to identify changes in signalling pathway activity with latent program activity.

### Cell communication inference

The R package nichenetr (NicheNet; v1.0.0)(Browaeys et al., 2020) was used to infer cell communication within the tumour microenvironment that could contribute to a sample’s specific EMP program. Given the diversity of EMP programs, this was performed on all 160 tumours independently. For each sample, cancer cells annotated by ACTIONet as the EMP-associated archetype were defined as the “receiver” population and all cell types were considered as potential “sender” cells. The “gene set of interest” was defined as that sample’s specific EMP program and expressed genes were defined as those with a detection rate of at least 5%. For each sample, we considered top ligands as the top 20 ligands inferred to promote expression of the EMP-associated genes.

### Assessing effects of small molecule inhibitors on EMP

ScRNA-seq data from epithelial cancer cell lines in control and drug-treated culture conditions were acquired from McFarland et al. (McFarland et al., 2020) and Cook &Vanderhyden (Cook and Vanderhyden, 2020). Initial data processing was performed identically to tumour samples and cell lines with fewer than 100 measured cells were removed. EMP programs were defined for each cell line using ACTIONet with only data from control cultured cells. Genes associated with each program were defined and used as a sample-specific EMP gene set for scoring both control and drug-treated cells. We then used a linear model to compare the effects of MEK and TGFBR inhibition on EMP in each line.

## AUTHOR CONTRIBUTIONS

D.P.C. and B.C.V. conceived the study and wrote the manuscript. D.P.C. performed all analysis.

## ACKNOWLEDGEMENTS

We extend deep gratitude to the many developers and maintainers of open source software that keep this field running. We thank Pascale Robineau-Charette for helpful discussions and feedback. David P. Cook was supported by a CIHR Frederick Banting and Charles Best Canada Graduate Scholarships Doctoral Award.

---

**Figure S1.**
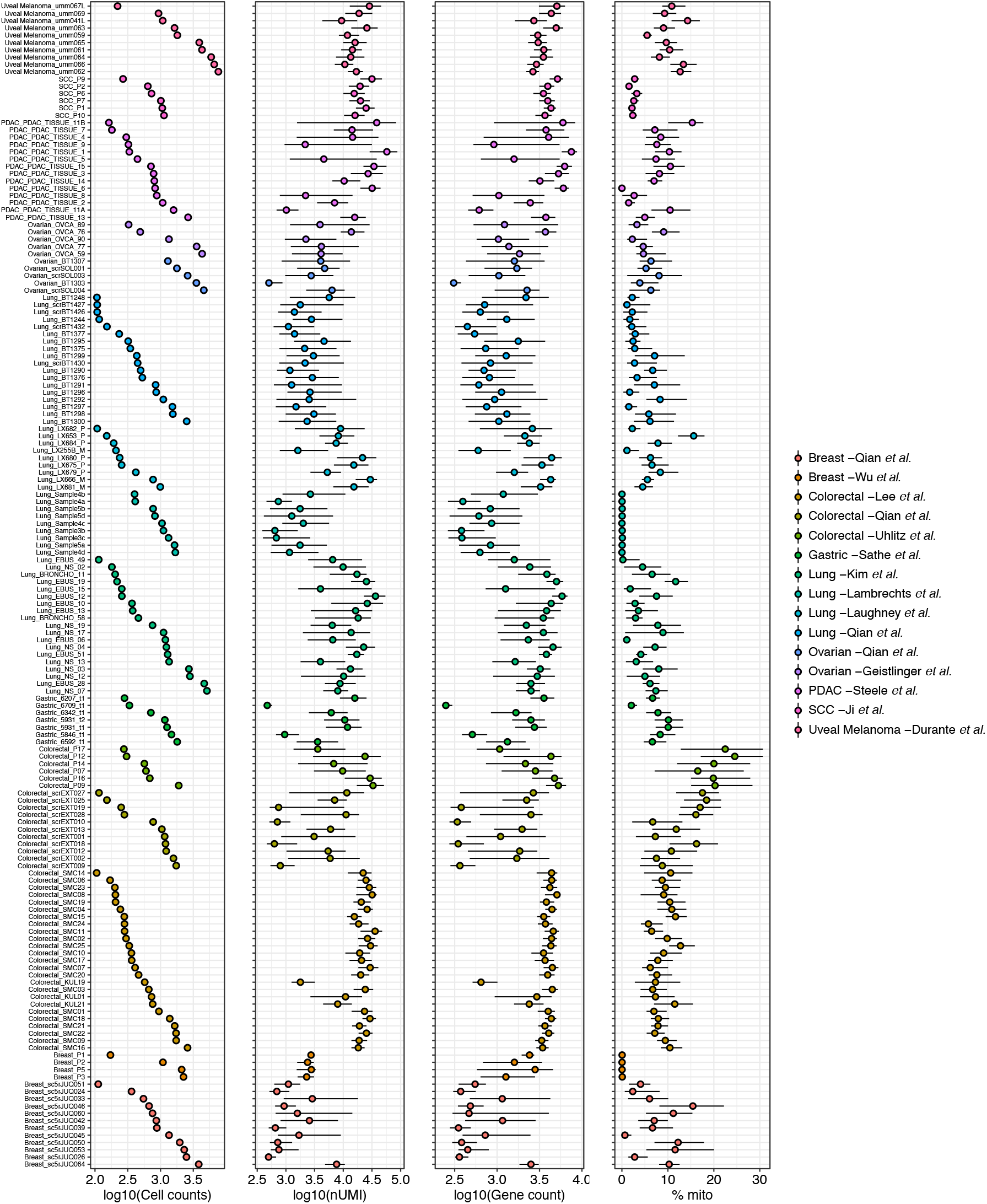
Quality control metrics for scRNA-seq data of malignant cells from 160 tumours. Plots showing the total number of cells, the distributions of unique transcript counts per cell, numbers of genes per cell, and percentage of mitochondrial reads per cell, respectively. Points represent the median value and the line spans the 25th-75th percentile.

**Figure S2.**
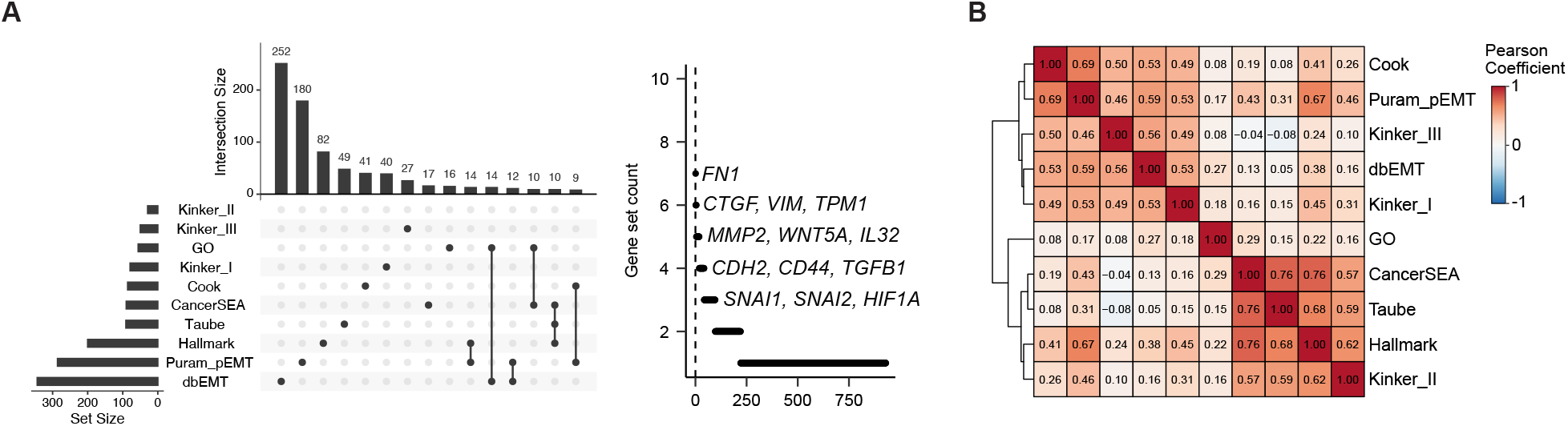
EMT gene sets are highly variable. **(A)** UpSet plot showing the overlap of gene composition for 10 EMT gene sets (left) and plot showing the number of gene sets within which each gene occurs (right). **(B)** Pearson correlation coefficients of gene set scores for each of the 10 EMT gene sets across the 160 tumours analyzed.

**Figure S3.**
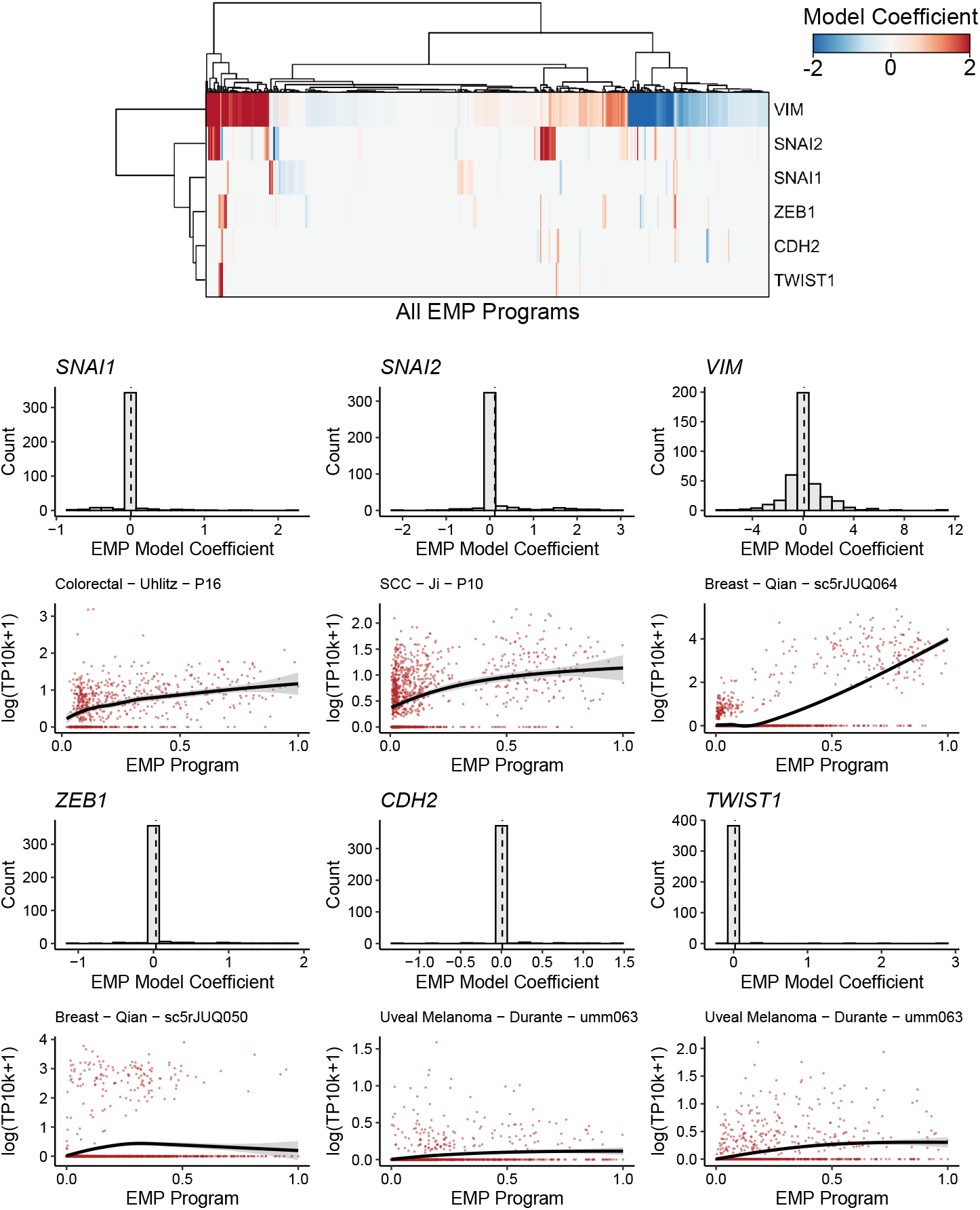
Association of canonical EMT genes in EMP programs. Top: Heatmap of EMP model coefficients for VIM, SNAI1, SNAI2, ZEB1, CDH2, and TWIST1. Bottom: Distribution of EMP model coefficients for each gene and expression values for programs with the highest coefficient for a given gene.

**Figure S4.**
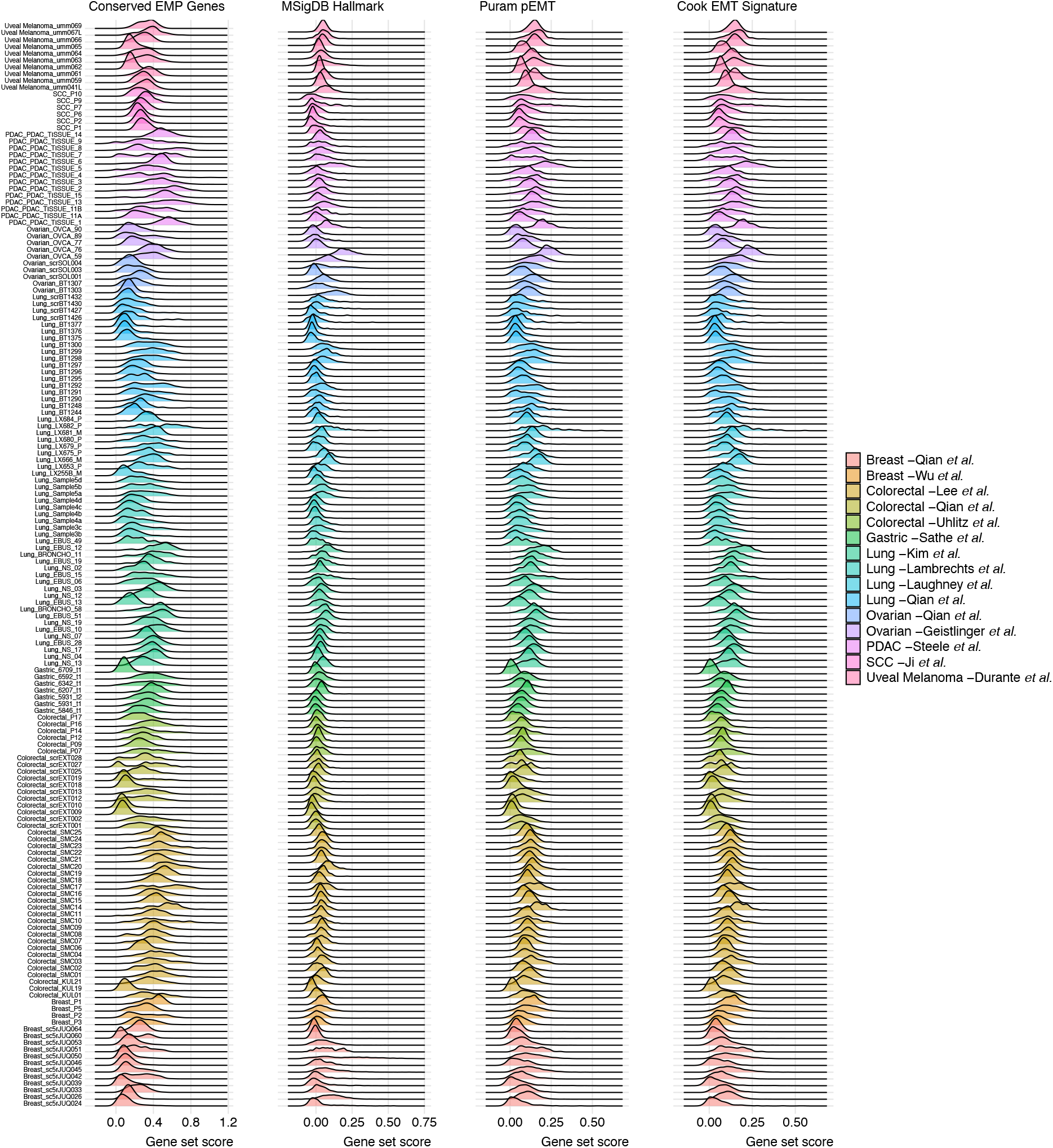
Distribution of EMP gene set scores. Density plots showing the distribution of gene set scores for the identified EMP signature (left) and three public EMT gene sets in malignant cells from all 160 tumours.

**Figure S5.**
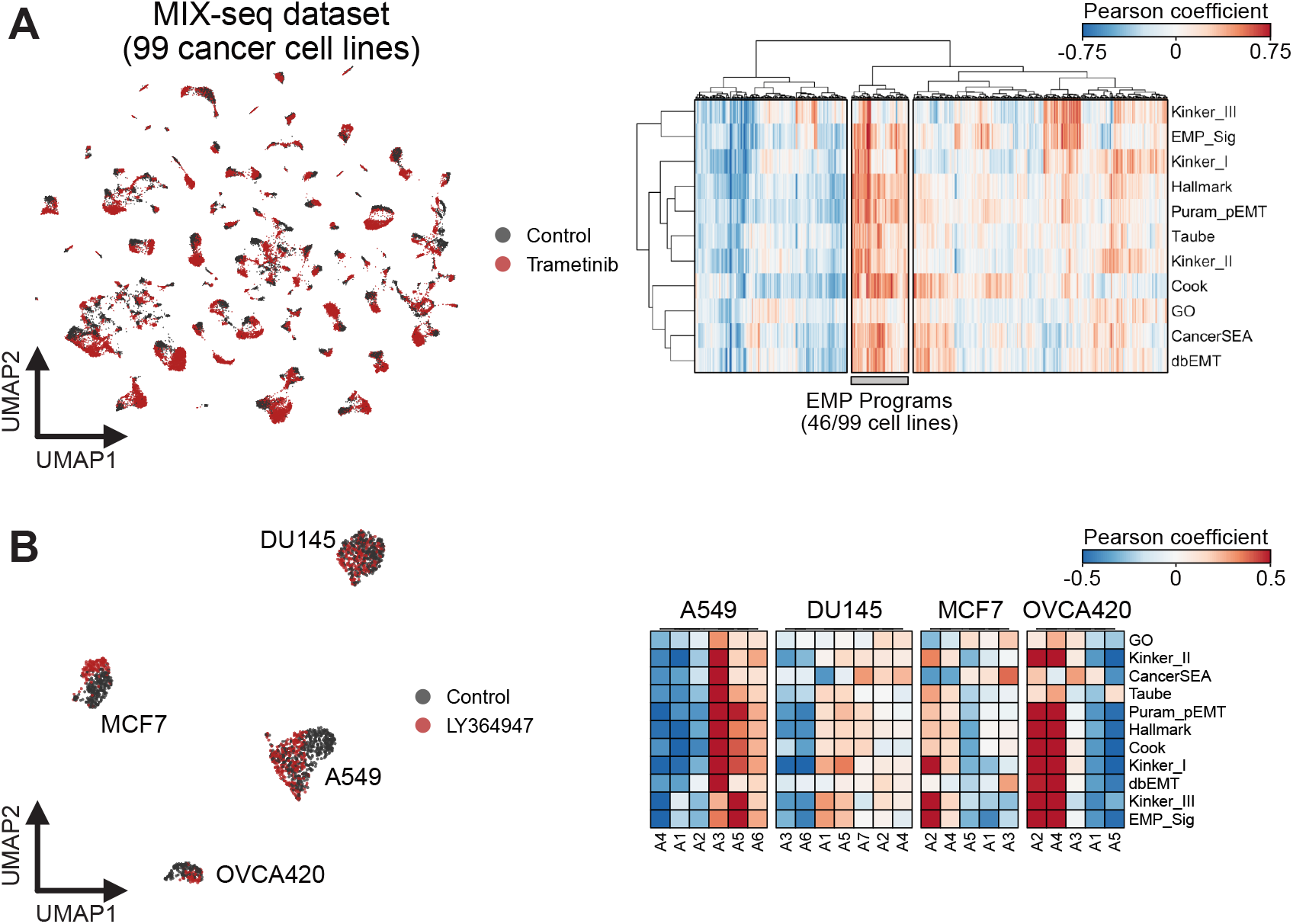
Identifying EMP signatures in cancer cell lines. **(A)** UMAP embedding of 99 cancer cell lines from the MIX-seq data set (left) and pearson correlation coefficients of archetype activity scores and EMT gene set scores for all cells. **(B)** Same as (A), but for scRNA-seq data of A549, DU145, MCF7, and OVCA420 cells from Cook &Vanderhyden, 2020.

## REFERENCES

Aiello, N.M., Brabletz, T., Kang, Y., Nieto, M.A., Weinberg, R.A., and Stanger, B.Z. (2017). Upholding a role for EMT in pancreatic cancer metastasis. Nature 547, E7–E8.

Aiello, N.M., Maddipati, R., Norgard, R.J., Balli, D., Li, J., Yuan, S., Yamazoe, T., Black, T., Sahmoud, A., Furth, E.E., et al. (2018). EMT Subtype Influences Epithelial Plasticity and Mode of Cell Migration. Dev. Cell 45, 681–695.e4.

Aran, D., Sirota, M., and Butte, A.J. (2015). Systematic pan-cancer analysis of tumour purity. Nat. Commun. 6, 8971.

Bhatia, S., Wang, P., Toh, A., and Thompson, E.W. (2020). New Insights Into the Role of Phenotypic Plasticity and EMT in Driving Cancer Progression. Front Mol Biosci 7, 71.

Bierie, B., Pierce, S.E., Kroeger, C., Stover, D.G., Pattabiraman, D.R., Thiru, P., Liu Donaher, J., Reinhardt, F., Chaffer, C.L., Keckesova, Z., et al. (2017). Integrin-β4 identifies cancer stem cell-enriched populations of partially mesenchymal carcinoma cells. Proc. Natl. Acad. Sci. U. S. A. 114, E2337–E2346.

Browaeys, R., Saelens, W., and Saeys, Y. (2020). NicheNet: modeling intercellular communication by linking ligands to target genes. Nat. Methods 17, 159–162.

Cook, D.P., and Vanderhyden, B.C. (2020). Context specificity of the EMT transcriptional response. Nat. Commun. 11, 2142.

Dongre, A., and Weinberg, R.A. (2019). New insights into the mechanisms of epithelial-mesenchymal transition and implications for cancer. Nat. Rev. Mol. Cell Biol. 20, 69–84.

Durante, M.A., Rodriguez, D.A., Kurtenbach, S., Kuznetsov, J.N., Sanchez, M.I., Decatur, C.L., Snyder, H., Feun, L.G., Livingstone, A.S., and Harbour, J.W. (2020). Single-cell analysis reveals new evolutionary complexity in uveal melanoma. Nat. Commun. 11, 496.

Fischer, K.R., Durrans, A., Lee, S., Sheng, J., Li, F., Wong, S.T.C., Choi, H., El Rayes, T., Ryu, S., Troeger, J., et al. (2015). Epithelial-to-mesenchymal transition is not required for lung metastasis but contributes to chemoresistance. Nature 527, 472–476.

Fischer, K.R., Altorki, N.K., Mittal, V., and Gao, D. (2017). Fischer et al. reply. Nature 547, E5–E6.

Gabbert, H., Wagner, R., Moll, R., and Gerharz, C.D. (1985). Tumor dedifferentiation: an important step in tumor invasion. Clin. Exp. Metastasis 3, 257–279.

Ganesh, K., Basnet, H., Kaygusuz, Y., Laughney, A.M., He, L., Sharma, R., O’Rourke, K.P., Reuter, V.P., Huang, Y.-H., Turkekul, M., et al. (2020). L1CAM defines the regenerative origin of metastasis-initiating cells in colorectal cancer. Nat Cancer 1, 28–45.

Geistlinger, L., Oh, S., Ramos, M., Schiffer, L., LaRue, R.S., Henzler, C.M., Munro, S.A., Daughters, C., Nelson, A.C., Winterhoff, B.J., et al. (2020). Multiomic Analysis of Subtype Evolution and Heterogeneity in High-Grade Serous Ovarian Carcinoma. Cancer Res. 80, 4335–4345.

Hafemeister, C., and Sat?a, R. (2019). Normalization and variance stabilization of single-cell RNA-seq data using regularized negative binomial regression. Genome Biol. 20, 296.

Hao, Y., Hao, S., Andersen-Nissen, E., Mauck, W.M., Zheng, S., Butler, A., Lee, M.J., Wilk, A.J., Darby, C., Zagar, M., et al. (2020). Integrated analysis of multimodal single-cell data.

Hoadley, K.A., Yau, C., Hinoue, T., Wolf, D.M., Lazar, A.J., Drill, E., Shen, R., Taylor, A.M., Cherniack, A.D., Thorsson, V., et al. (2018). Cell-of-Origin Patterns Dominate the Molecular Classification of 10,000 Tumors from 33 Types of Cancer. Cell 173, 291–304.e6.

Holland, C.H., Szalai, B., and Saez-Rodriguez, J. (2020a). Transfer of regulatory knowledge from human to mouse for functional genomics analysis. Biochim. Biophys. Acta Gene Regul. Mech. 1863, 194431.

Holland, C.H., Tanevski, J., Perales-Patón, J., Gleixner, J., Kumar, M.P., Mereu, E., Joughin, B.A., Stegle, O., Lauffenburger, D.A., Heyn, H., et al. (2020b). Robustness and applicability of transcription factor and pathway analysis tools on single-cell RNA-seq data. Genome Biol. 21, 36.

Horn, L.A., Fousek, K., and Palena, C. (2020). Tumor Plasticity and Resistance to Immunotherapy. Trends Cancer Res. 6, 432–441.

Ibrahim, M.M., and Kramann, R. (2019). genesorteR: Feature Ranking in Clustered Single Cell Data.

Isella, C., Terrasi, A., Bellomo, S.E., Petti, C., Galatola, G., Muratore, A., Mellano, A., Senetta, R., Cassenti, A., Sonetto, C., et al. (2015). Stromal contribution to the colorectal cancer transcriptome. Nat. Genet. 47, 312–319.

Izar, B., Tirosh, I., Stover, E.H., Wakiro, I., Cuoco, M.S., Alter, I., Rodman, C., Leeson, R., Su, M.-J., Shah, P., et al. (2020). A single-cell landscape of high-grade serous ovarian cancer. Nat. Med. 26, 1271–1279.

Ji, A.L., Rubin, A.J., Thrane, K., Jiang, S., Reynolds, D.L., Meyers, R.M., Guo, M.G., George, B.M., Mollbrink, A., Bergenstråhle, J., et al. (2020). Multimodal Analysis of Composition and Spatial Architecture in Human Squamous Cell Carcinoma. Cell 182, 1661–1662.

Jia, D., Jolly, M.K., Tripathi, S.C., Den Hollander, P., Huang, B., Lu, M., Celiktas, M., Ramirez-Peña, E., Ben-Jacob, E., Onuchic, J.N., et al. (2017). Distinguishing mechanisms underlying EMT tristability. Cancer Converg 1, 2.

Kim, N., Kim, H.K., Lee, K., Hong, Y., Cho, J.H., Choi, J.W., Lee, J.-I., Suh, Y.-L., Ku, B.M., Eum, H.H., et al. (2020). Single-cell RNA sequencing demonstrates the molecular and cellular reprogramming of metastatic lung adenocarcinoma. Nat. Commun. 11, 2285.

Kinker, G.S., Greenwald, A.C., Tal, R., Orlova, Z., Cuoco, M.S., McFarland, J.M., Warren, A., Rodman, C., Roth, J.A., Bender, S.A., et al. (2020). Pan-cancer single-cell RNA-seq identifies recurring programs of cellular heterogeneity. Nat. Genet. 52, 1208–1218.

Klein, A.M., Mazutis, L., Akartuna, I., Tallapragada, N., Veres, A., Li, V., Peshkin, L., Weitz, D.A., and Kirschner, M.W. (2015). Droplet barcoding for single-cell transcriptomics applied to embryonic stem cells. Cell 161, 1187–1201.

Kröger, C., Afeyan, A., Mraz, J., Eaton, E.N., Reinhardt, F., Khodor, Y.L., Thiru, P., Bierie, B., Ye, X., Burge, C.B., et al. (2019). Acquisition of a hybrid E/M state is essential for tumorigenicity of basal breast cancer cells. Proc. Natl. Acad. Sci. U. S. A. 116, 7353–7362.

Lambrechts, D., Wauters, E., Boeckx, B., Aibar, S., Nittner, D., Burton, O., Bassez, A., Decaluwé, H., Pircher, A., Van den Eynde, K., et al. (2018). Phenotype molding of stromal cells in the lung tumor microenvironment. Nat. Med. 24, 1277–1289.

Laughney, A.M., Hu, J., Campbell, N.R., Bakhoum, S.F., Setty, M., Lavallée, V.-P., Xie, Y., Masilionis, I., Carr, A.J., Kottapalli, S., et al. (2020). Regenerative lineages and immune-mediated pruning in lung cancer metastasis. Nat. Med. 26, 259–269.

Lee, H.-O., Hong, Y., Etlioglu, H.E., Cho, Y.B., Pomella, V., Van den Bosch, B., Vanhecke, J., Verbandt, S., Hong, H., Min, J.-W., et al. (2020). Lineage-dependent gene expression programs influence the immune landscape of colorectal cancer. Nat. Genet. 52, 594–603.

Liberzon, A., Birger, C., Thorvaldsdóttir, H., Ghandi, M., Mesirov, J.P., and Tamayo, P. (2015). The Molecular Signatures Database (MSigDB) hallmark gene set collection. Cell Syst 1, 417–425.

Macosko, E.Z., Basu, A., Sat?a, R., Nemesh, J., Shekhar, K., Goldman, M., Tirosh, I., Bialas, A.R., Kamitaki, N., Martersteck, E.M., et al. (2015). Highly Parallel Genome-wide Expression Profiling of Individual Cells Using Nanoliter Droplets. Cell 161, 1202–1214.

Marjanovic, N.D., Hofree, M., Chan, J.E., Canner, D., Wu, K., Trakala, M., Hartmann, G.G., Smith, O.C., Kim, J.Y., Evans, K.V., et al. (2020). Emergence of a High-Plasticity Cell State during Lung Cancer Evolution. Cancer Cell 38, 229–246.e13.

McFaline-Figueroa, J.L., Hill, A.J., Qiu, X., Jackson, D., Shendure, J., and Trapnell, C. (2019). A pooled single-cell genetic screen identifies regulatory checkpoints in the continuum of the epithelial-to-mesenchymal transition. Nat. Genet. 51, 1389–1398.

McFarland, J.M., Paolella, B.R., Warren, A., Geiger-Schuller, K., Shibue, T., Rothberg, M., Kuksenko, O., Colgan, W.N., Jones, A., Chambers, E., et al. (2020). Multiplexed single-cell transcriptional response profiling to define cancer vulnerabilities and therapeutic mechanism of action. Nat. Commun. 11, 4296.

Mohammadi, S., Davila-Velderrain, J., and Kellis, M. (2020). A multiresolution framework to characterize single-cell state landscapes. Nat. Commun. 11, 5399.

Nieto, M.A., Huang, R.Y.-J., Jackson, R.A., and Thiery, J.P. (2016). EMT: 2016. Cell 166, 21–45.

Pastushenko, I., and Blanpain, C. (2019). EMT Transition States during Tumor Progression and Metastasis. Trends Cell Biol. 29, 212–226.

Pastushenko, I., Brisebarre, A., Sifrim, A., Fioramonti, M., Revenco, T., Boumahdi, S., Van Keymeulen, A., Brown, D., Moers, V., Lemaire, S., et al. (2018). Identification of the tumour transition states occurring during EMT. Nature 556, 463–468.

Peixoto, P., Etcheverry, A., Aubry, M., Missey, A., Lachat, C., Perrard, J., Hendrick, E., Delage-Mourroux, R., Mosser, J., Borg, C., et al. (2019). EMT is associated with an epigenetic signature of ECM remodeling genes. Cell Death Dis. 10, 205.

Puram, S.V., Tirosh, I., Parikh, A.S., Patel, A.P., Yizhak, K., Gillespie, S., Rodman, C., Luo, C.L., Mroz, E.A., Emerick, K.S., et al. (2017). Single-Cell Transcriptomic Analysis of Primary and Metastatic Tumor Ecosystems in Head and Neck Cancer. Cell 171, 1611–1624.e24.

Qian, J., Olbrecht, S., Boeckx, B., Vos, H., Laoui, D., Etlioglu, E., Wauters, E., Pomella, V., Verbandt, S., Busschaert, P., et al. (2020). A pan-cancer blueprint of the heterogeneous tumor microenvironment revealed by single-cell profiling. Cell Res. 30, 745–762.

Raghavan, S., Winter, P.S., Navia, A.W., Williams, H.L., DenAdel, A., Kalekar, R.L., Galvez-Reyes, J., Lowder, K.E., Mulugeta, N., Raghavan, M.S., et al. Transcriptional subtype-specific microenvironmental crosstalk and tumor cell plasticity in metastatic pancreatic cancer.

Ramesh, V., Brabletz, T., and Ceppi, P. (2020). Targeting EMT in Cancer with Repurposed Metabolic Inhibitors. Trends Cancer Res. 6, 942– 950.

Sathe, A., Grimes, S.M., Lau, B.T., Chen, J., Suarez, C., Huang, R.J., Poultsides, G., and Ji, H.P. (2020). Single-Cell Genomic Characterization Reveals the Cellular Reprogramming of the Gastric Tumor Microenvironment. Clin. Cancer Res. 26, 2640–2653.

Schubert, M., Klinger, B., Klünemann, M., Sieber, A., Uhlitz, F., Sauer, S., Garnett, M.J., Blüthgen, N., and Saez-Rodriguez, J. (2018). Perturbation-response genes reveal signaling footprints in cancer gene expression. Nat. Commun. 9, 20.

Sharma, A., Cao, E.Y., Kumar, V., Zhang, X., Leong, H.S., Wong, A.M.L., Ramakrishnan, N., Hakimullah, M., Teo, H.M.V., Chong, F.T., et al. (2018). Longitudinal single-cell RNA sequencing of patient-derived primary cells reveals drug-induced infidelity in stem cell hierarchy. Nature Communications 9.

Steele, N.G., Carpenter, E.S., Kemp, S.B., Sirihorachai, V.R., Stephanie The, Delrosario, L., Lazarus, J., Amir, E.-A.D., Gunchick, V., Espinoza, C., et al. (2020). Multimodal mapping of the tumor and peripheral blood immune landscape in human pancreatic cancer. Nature Cancer 1, 1097–1112.

Stemmler, M.P., Eccles, R.L., Brabletz, S., and Brabletz, T. (2019). Non-redundant functions of EMT transcription factors. Nat. Cell Biol. 21, 102–112.

Taube, J.H., Herschkowitz, J.I., Komurov, K., Zhou, A.Y., Gupta, S., Yang, J., Hartwell, K., Onder, T.T., Gupta, P.B., Evans, K.W., et al. (2010). Core epithelial-to-mesenchymal transition interactome gene-expression signature is associated with claudin-low and metaplastic breast cancer subtypes. Proc. Natl. Acad. Sci. U. S. A. 107, 15449–15454.

Thorsson, V., Gibbs, D.L., Brown, S.D., Wolf, D., Bortone, D.S., Ou Yang T.-H., Porta-Pardo, E., Gao, G.F., Plaisier, C.L., Eddy, J.A., et al. (2019). The Immune Landscape of Cancer. Immunity 51, 411–412.

Uhlitz, F., Bischoff, P., Peidli, S., Sieber, A., Obermayer, B., Blanc, E., Trinks, A., Lüthen, M., Ruchiy, Y., Sell, T., et al. Mitogen-activated protein kinase activity drives cell trajectories in colorectal cancer.

Williams, E.D., Gao, D., Redfern, A., and Thompson, E.W. (2019). Controversies around epithelial-mesenchymal plasticity in cancer metastasis. Nat. Rev. Cancer 19, 716–732.

Yang, J., Antin, P., Berx, G., Blanpain, C., Brabletz, T., Bronner, M., Campbell, K., Cano, A., Casanova, J., Christofori, G., et al. (2020). Guidelines and definitions for research on epithelial-mesenchymal transition. Nat. Rev. Mol. Cell Biol. 21, 341–352.

Ye, X., Brabletz, T., Kang, Y., Longmore, G.D., Nieto, M.A., Stanger, B.Z., Yang, J., and Weinberg, R.A. (2017). Upholding a role for EMT in breast cancer metastasis. Nature 547, E1–E3.

Yuan, H., Yan, M., Zhang, G., Liu, W., Deng, C., Liao, G., Xu, L., Luo, T., Yan, H., Long, Z., et al. (2019). CancerSEA: a cancer single-cell state atlas. Nucleic Acids Res. 47, D900–D908.

Zadran, S., Arumugam, R., Herschman, H., Phelps, M.E., and Levine, R.D. (2014). Surprisal analysis characterizes the free energy time course of cancer cells undergoing epithelial-to-mesenchymal transition. Proc. Natl. Acad. Sci. U. S. A. 111, 13235–13240.

Zhao, M., Kong, L., Liu, Y., and Qu, H. (2015). dbEMT: an epithelial-mesenchymal transition associated gene resource. Sci. Rep. 5, 11459.

Zheng, X., Carstens, J.L., Kim, J., Scheible, M., Kaye, J., Sugimoto, H., Wu, C.-C., LeBleu, V.S., and Kalluri, R. (2015). Epithelial-to-mesenchymal transition is dispensable for metastasis but induces chemoresistance in pancreatic cancer. Nature 527, 525–530.

